# Effective Biophysical Modeling of Cell Free Transcription and Translation Processes

**DOI:** 10.1101/2020.02.25.964841

**Authors:** Abhinav Adhikari, Michael Vilkhovoy, Sandra Vadhin, Ha Eun Lim, Jeffrey D. Varner

**Affiliations:** Robert Frederick Smith School of Chemical and Biomolecular Engineering College of Engineering, Cornell University, Ithaca, NY 14853

## Abstract

Transcription and translation are at the heart of metabolism and signal transduction. In this study, we developed an effective biophysical modeling approach to simulate transcription and translation processes. We tested this approach by simulating the dynamics of two cell free synthetic circuits. First, we considered a simple circuit in which sigma factor 70 induced the expression of green fluorescent protein. This relatively simple case was then followed by a more complex negative feedback circuit in which two control genes were coupled to the expression of a third reporter gene, green fluorescent protein. While many of the model parameters were estimated from previous biophysical literature, the remaining unknown model parameters for each circuit were estimated from messenger RNA (mRNA) and protein measurements using multi-objective optimization. In particular, either the literature parameter estimates were used directly in the model simulations, or characteristic literature values were used to establish feasible ranges for the multiobjective parameter search. Next, global sensitivity analysis was used to determine the influence of individual model parameters on the expression dynamics. Taken together, the effective biophysical modeling approach captured the expression dynamics, including the transcription dynamics, for the two synthetic cell free circuits. While we considered only two circuits here, this approach could potentially be extended to simulate other genetic circuits in both cell free and whole cell biomolecular applications. The model code, parameters, and analysis scripts are available for download under an MIT software license from the Varnerlab GitHub repository.

## Introduction

Transcription (TX) and translation (TL), the processes by which information stored in DNA is converted to a working protein, are at the center of metabolism and signal transduction programs important to biotechnology and human heath. For example, evolutionarily conserved developmental programs such as the epithelial to mesenchymal transition (EMT) [1], or retinoic acid induced differentiation [2], rely on multiple rounds of highly coordinated gene expression. From the perspective of biotechnology, even relatively simple industrially important organisms such as *Escherichia coli,* have intricate regulatory networks which control the metabolic state of the cell in response to changing nutrient conditions [3, 4]. Understanding the dynamics of regulatory networks can be greatly facilitated by mathematical models. A majority of these models fall into three categories: logical, continuous, and stochastic models [5]. Logical models such as Boolean networks [6] developed using a variety of approaches and data [7] represent the state of each network entity as a discrete variable, provide a quick but qualitative description of the behavior of the regulatory network. Linear and non-linear ordinary differential equation (ODE) models fall into the second category, and they generally provide a detailed picture of the network dynamics, although they can be non-physical models, e.g., relying on a gene signal perspective [8]. Lastly, stochastic models describe the interactions between individual molecules, and discrete reaction events [9–12]. Model choice depends on criteria such as speed, the level of detail required and the quantity of experimental data available to estimate the model parameters. While the end goal of the models might be to accurately predict *in vivo* behavior, living systems do not necessarily provide an ideal experimental platform. For example, although there have been significant advancements in metabolomics e.g., [13], the rigorous quantification of intracellular messenger RNA (mRNA) copy number or protein abundance remains challenging. Toward this challenge, cell free systems offer several advantages for the study of transcription and translation processes.

Cell free biology has historically been an important tool to study the fundamental biological mechanisms involved with gene expression. In the 1950s, cell free systems were used to explore the incorporation of amino acids into proteins [14–16], and the role of adenosine triphosphate (ATP) in protein production [17]. Further, *E. coli* extracts were used by Nirenberg and Matthaei in 1961 to demonstrate templated translation [18, 19], leading to a Nobel Prize in 1968 for deciphering the codon code. More recently, as advancements in extract preparation and energy regeneration have extend their durability, the usage of cell free systems has also expanded to both small- and large-scale biotechnology and biomanufacturing applications [20, 21]. Today, cell free systems have been implemented for therapeutic protein and vaccine production [22–24], biosensing [25], genetic part prototyping [26] and minimal cell systems [27]. The versatility of cell free systems offers a tremendous opportunity for the systems-level experimental and computational study of biological mechanism. For example, a number of ordinary differential equation based cell free models have been developed [28–32]. However, despite the obvious advantages offered by a cell free system, experimental determination of the kinetic parameters for these models is often not feasible. For instance, the cell free modeling study of Horvath and coworkers (which included a description of transcription and translation, and the underlying metabolism supplying energy and precursors for transcription and translation), had over 800 unknown model parameters [33]. Moreover, the construction, identification and validation of the Horvath model took well over a year to complete. Thus, constructing, identifying and validating biophysically motivated cell free models, which are simultaneously manageable, is a key challenge. Toward this challenge, effective modeling approaches which coarse grain biological details but remain firmly rooted in a biophysical perspective, could be an important tool.

In this study, we developed an effective biophysical modeling approach to simulate cell free transcription and translation processes. We tested this approach by simulating the dynamics of two cell free synthetic circuits. First, we considered a simple circuit 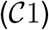 in which endogenous sigma factor 70 (σ_70_) induced the expression of a fast maturing dual emission green fluorescent protein variant (deGFP). This relatively simple case was then followed by a more complex negative feedback circuit 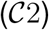 where two control genes were coupled to the expression of deGFP. Characteristic values for many of the model parameters were estimated from previous biophysical literature, while the remaining unknown model parameters for each circuit were estimated from messenger RNA (mRNA) and protein measurements using multiobjective optimization. In particular, either the literature parameter estimates were used directly in the model simulations, or characteristic literature values were used to establish feasible ranges for the multiobjective parameter search. Next, Morris sensitivity analysis was used to determine the influence of individual model parameters on the expression dynamics of each circuit. For 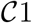, the sensitivity results were informative, but expected. However, for 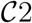, the analysis hierarchically stratified the parameters (and associated model species) into local versus global categories. For example, parameters that controlled the abundance of lambda phage repressor protein (cI-ssrA), a master circuit regulator in 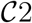, were globally important as they influenced all other species. On the other hand, the parameters that influenced deGFP levels (the endpoint of both circuits) were only locally important to deGFP. Taken together, the effective biophysical modeling approach captured the expression dynamics, including the transcription dynamics, for two synthetic cell free circuits. While we considered only two circuits here, this approach could potentially be extended to simulate other genetic circuits in both cell free and whole cell biomolecular applications. The model code, parameters, and analysis scripts are available under an MIT software license from the Varnerlab GitHub repository [34].

## Materials and Methods

### myTX/TL cell-free protein synthesis

The cell-free protein synthesis (CFPS) reactions were carried out using the myTXTL Sigma 70 Master Mix (Arbor Biosciences) in 1.5 mL Eppendorf tubes. The working volume of all the reactions was 12 *μ*L, composed of the Sigma 70 Master Mix (9 *μ*L) and the plasmids (3 *μ*L total): P70a-deGFP (5 nM) for the single-gene system; P70a-deGFP-ssrA (8 nM), P70a-S28 (1.5 nM) and P28a-cI-ssrA (1 nM) for the negative feedback circuit. The plasmids were bought in lyophilized form (Arbor Biosciences) and purified using QIAprep Spin Miniprep Kit (Qiagen) using cell lines DH5-Alpha (for P28a-cI-ssrA) or KL740 (for P70a-deGFP, P70a-deGFP-ssrA, and P70a-S28). The CFPS reactions were incubated at 29°C.

### mRNA and protein quantification

Following each CFPS run, the total RNA was extracted from 1 *μ*L of the reaction mixture using PureLink RNA Mini Kit (Thermo Fisher Scientific) and stored at −80°C. The quantitative RT-PCR reactions were done using Applied Biosystems™ TaqMan™ RNA-to-CT™ 1-Step Kit and Custom TaqMan Gene Expression Assays (Thermo Fisher Scientific). The standard curve method was used to determine absolute mRNA concentrations for the samples. At least three technical replicates were performed for each standard. The mRNA standards were prepared as follows: separate CFPS reactions for 5 nM of plasmids (P70a-S28, P70a-deGFP, and P70a-deGFP-ssrA) were carried out for 2 hours. Total RNA was extracted using the full reaction volume. This was followed by the removal of 16S and 23S rRNA using the MICROBExpress™ Bacterial mRNA Enrichment Kit (Life Technologies Corporation). Lastly, the MEGAclear™ Kit (Life Technologies Corporation) was used to further purify the mRNA. The mRNA concentrations were determined using the Qubit™ RNA assay kit (ThermoFisher Scientific). The concentration of cI-ssrA mRNA was quantified using the deGFP-ssrA mRNA standard. Green fluorescent protein (deGFP) fluorescence was measured using the Varioskan Lux plate reader at 488 nm (excitation) and 535 nm (emission). At the end of the CFPS run, 3 *μ*L of the reaction mixture was diluted in 33 *μ*L phosphate buffered saline (PBS) and stored at −80°C. The fluorescence was measured in triplicate with 10 *μ*L each of this mixture. For all measurements, at least three biological replicates were performed.

### Formulation and solution of model equations

Consider a gene regulatory motif with genes 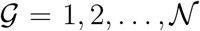. Each node in this motif is governed by two differential equations, one for mRNA (*m,_j_*) and the corresponding protein (*p_j_*):

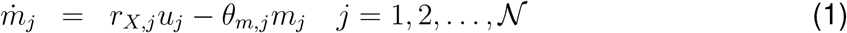

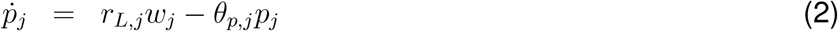

The term *r_X,j_u_j_* in the mRNA balance, which denotes the rate of transcription for gene j, is the product of a kinetic limit *r_X,j_* (nmol h^-1^) and a transcription control function 0 ≤ *u_j_* (…) ≤ 1 (dimensionless). Similarly, the rate of translation of mRNA j, denoted by *r_L,j_w_j_*, is also the product of the kinetic limit of translation and a translational control term 0 ≤ *w_j_* (…) ≤ 1 (dimensionless). Lastly, *θ*_*,*j*_ denotes the first-order rate constant (h^-1^) governing degradation of protein and mRNA. The model equations, encoded in the Julia programming language [35], were automatically generated using the JuGRN tool [36]. The model equations were solved numerically using the `Rosenbrock23` routine of the `DifferentialEquations.jl` package [37].

### Derivation of the transcription and translation kinetic limits

The key idea behind the derivation of the kinetic limit of transcription and translation is that the polymerase (or ribosome) acts as a pseudo-enzyme; it binds a substrate (gene or message), reads the gene (or message), and then dissociates. Thus, we might expect that we could use a strategy similar to enzyme kinetics to derive an expression for *r_X,j_* (or *r_L,j_*). Toward this idea, the kinetic limit of transcription *r_X,j_* was developed by proposing a simple four elementary step model:

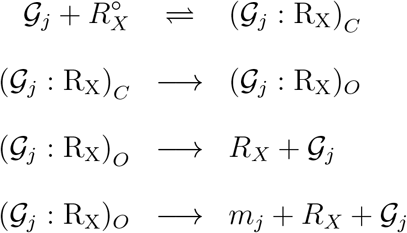

where 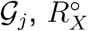 denote the gene and *free* RNAP concentration, and 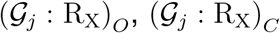 denote the open and closed complex concentrations, respectively. Let the kinetic limit of transcription be directly proportional to the concentration of the open complex:

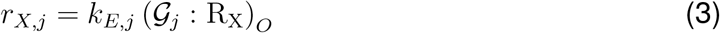

where *k_E,j_* is the elongation rate constant for gene *j*. Then the functional form for *r_X,j_* can be derived:

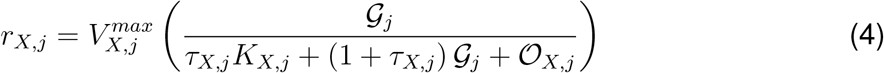

where 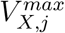 denotes the maximum transcription rate (nM/h) of gene *j* and:

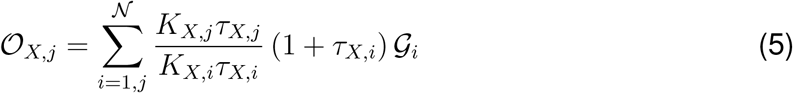

denotes the coupling of the transcription of gene *j* with the other genes in the system through competition for RNA polymerase. In a similar way, we developed a model for the translational kinetic limit:

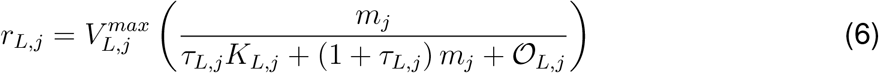

where 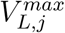 denotes the maximum translation rate (nM/h) and:

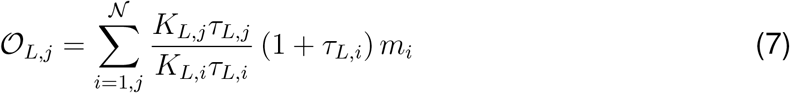

describes the coupling of the translation of mRNA *j* with other messages in the system because of kinetic competition for available ribosomes. The saturation and time constants for each case were defined as 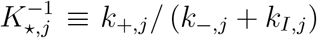 and 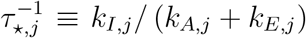, respectively. In this study, the 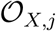 and 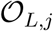 terms were neglected as both circuits had either only one, or a small number of genes.

The maximum transcription rate 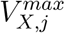 was formulated as:

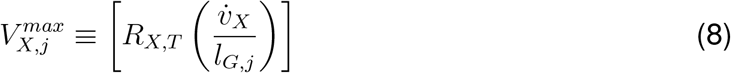

The term *R_X,T_* denotes the total RNA polymerase concentration (nM), 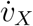 denotes the RNA polymerase elongation rate (nt/h), *l_G,j_* denotes the length of gene *j* in nucleotides (nt). Similarly, the maximum translation rate 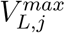 was formulated as:

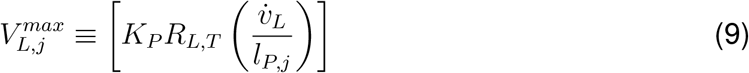

The term *R_L,T_* denotes the total ribosome pool, *K_P_* denotes the polysome amplification constant, 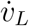 denotes the ribosome elongation rate (amino acids per hour), and *l_P,j_* denotes the length of protein *j* (aa).

*Control functions u and w*. Values of the control functions *u* (…) and *w* (…) describe the regulation of transcription and translation. Ackers et al., borrowed from statistical mechanics to recast the *u*(…) function as the probability that a system exists in a configuration which leads to expression [38]. The idea of recasting *u*(…) as the probability of expression was also developed (apparently independently) by Bailey and coworkers in a series of papers modeling the *lac* operon, see [39]. More recently, Moon and Voigt adapted a similar approach when modeling the expression of synthetic circuits in *E. coli* [40]. The *u*(…) function is formulated as:

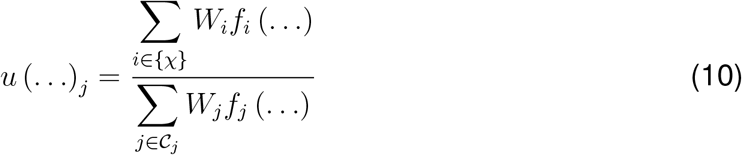

where *W_i_*, (dimensionless) denotes the weight of configuration *i*, while *f_i_* (…) (dimensionless) is a binding function (taken to be a hill-type function) describing the fraction of bound activator/inhibitor for configuration *i*. The summation in the numerator of Eq. (10) is over the set of configurations that lead to expression (denoted as*χ*), while the summation in the numerator is over the set of all possible configurations for gene *j* (denoted as 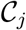). Thus, *u* (…)_*j*_ can be thought of as the fraction of all possible configurations that lead to expression. The weights *W*, are related to the Gibbs energy of configuration *i*: *W_i_*, = exp (-Δ*G_i_*/*RT*) where Δ*G_i_*, denotes the molar Gibbs energy for configuration *i* (kJ/mol), *R* denotes the ideal gas constant (kJ mol^-1^ K^-1^), and *T* denotes the system temperature (Kelvin) [38].

The translational control function *w* (…) described the loss of cell free translational activity. Loss of translational activity could be a function of many factors, including depletion of metabolic resources. However, in this study we assumed a translational activity factor *ϵ* of the form:

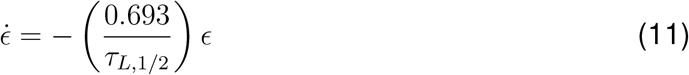

where *τ*_*L*,1/2_ denotes the half-life of translation (estimated from the experimental data) and *ϵ*(0) = 1. The translational control variable was then given by *w_i_*, = *ϵ* for all translation processes.

### Estimation of model parameters

Model parameters were estimated using the Pareto Optimal Ensemble Technique in the Julia programming language (JuPOETs) [41]. Several of the model parameters were estimated directly from literature, or were set by experimental conditions (Table 1). However, the remaining unknown parameters (k) were estimated from mRNA and protein concentration measurements using JuPOETs. JuPOETs is a multiobjective optimization approach which integrates simulated annealing with Pareto optimality to estimate parameter ensembles on or near the optimal tradeoff surface between competing training objectives. JuPOETs minimized training objectives of the form:

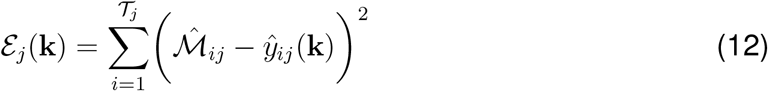

subject to the model equations, initial conditions, parameter bounds 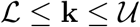 and maximum value constraints for some unmeasured protein species. The objective function(s) 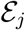 measured the squared difference between the model simulations and experimental measurements for training objective *j*. The symbol 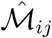 denotes an experimental observation at time index *i* for training objective *j*, while the symbol 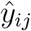 denotes the model simulation output at time index *i* from training objective *j*. The quantity *i* denotes the sampled time-index and 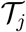 denotes the number of time points for experiment *j*. For the P70-deGFP model 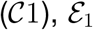 corresponded to mRNA deGFP, while 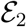 corresponded to the deGFP protein concentration. On the other hand, for the negative feedback model 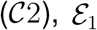 corresponded to mRNA deGFP-ssrA, 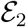 to mRNA *σ*_28_, 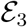 to mRNA cI-ssrA and 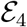 to the deGFP-ssrA protein concentration. Lastly, we penalized non-physical accumulation of the cI-ssrA protein (unmeasured) with a term of the form: 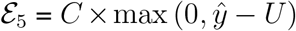 where *C* denotes a penalty parameter (*C* = 1 ×10^5^), 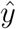 denotes the maximum cI-ssrA concentration, and *U* denotes a concentration upper bound (U = 100nM). For 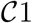, 11 parameters were estimated, while 33 parameters were estimated for 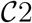.

**Table 1:** Characteristic parameters for TX/TL model equations. Key to references used in the table: (a) [45], (b) [66], (c) set by experiment, (d) [67], (e) estimated in this study, (f) [68], (g) [69], (h) calculated from plasmid sequence, (i) [47], (f) [53]

| Description | Parameter | Value | Units | Reference |
| --- | --- | --- | --- | --- |
| RNA polymerase concentration | $R_{X,T}$ | 0.06-0.075 | $\mu\text{M}$ | *a* |
| Ribosome concentration | $R_{L,T}$ | $< 2.3$ | $\mu\text{M}$ | *a,b* |
| $\sigma_{70}$ concentration | $\sigma_{70}$ | $< 35$ | nM | *a* |
| initial $\sigma_{28}$ concentration | $\sigma_{28}$ | $< 20$ | nM | *a* |
| Transcription elongation rate | $\dot{v}_X$ | 12-30 | nt/s | *a,d* |
| Translation elongation rate | $\dot{v}_L$ | 1-2 | aa/s | *a,b* |
| Transcription saturation coefficient | $K_X$ | 0.036 | $\mu\text{M}$ | *i* |
| Polysome amplification constant | $K_P$ | 10.0 | constant | *e* |
| Transcription initiation time | $k_I^X$ | 22 | s | *i* |
| Translation initiation time | $k_I^L$ | 1.5 | s | *e* |
| Default mRNA degradation coefficient | $\theta_m$ | 3.75 | $\text{h}^{-1}$ | *a* |
| Default protein degradation coefficient | $\theta_p$ | 0.462-1.89 | $\text{h}^{-1}$ | *f,g* |
| Gene concentration $\sigma_{28}$ | | 1.5 | nM | *c* |
| Gene concentration cl-ssrA |  | 1.0 | nM | *c* |
| Gene concentration deGFP-ssrA |  | 8.0 | nM | *c* |
| Gene length $\sigma_{28}$ | | 811 | nt | *h* |
| Gene length cl-ssrA |  | 850 | nt | *h* |
| Gene length deGFP-ssrA |  | 782 | nt | *h* |
| Protein length $\sigma_{28}$ | | 240 | aa | *h* |
| Protein length cl-ssrA |  | 248 | aa | *h* |
| Protein length deGFP-ssrA |  | 237 | aa | *h* |

Parameter bounds 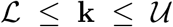 were established from literature, or from previous experimental or model analysis; parameter values estimated for the P70-deGFP model were also used to establish ranges for the negative feedback model. JuPOETs searched over Δ*G_i_, K_L_* and *τ_L,1/2_* values directly, while other unknown parameter values took the form of corrections to order of magnitude characteristic literature estimates. For example, we set the mRNA degradation rate constant (*θ_m_*) to a characteristic value taken from literature. Then, the degradation constant for any particular mRNA was represented as: *θ_m,i_* = *α_i_θ_m_*, where *α_i_*, was an unknown (but bounded) modifier. In this way, we guaranteed the parameter search (and the resulting estimated parameters) were within a specified range of literature values. We used this procedure for all degradation constants (both mRNA and protein) and all time constants (for both transcription and translation). The baseline parameter values are given in Table 1. JuPOETs was run for 20 generations for both models, and all parameters sets with Pareto rank less than or equal to two were collected for each generation. The JuPOETs parameter estimation routine is encoded in the `sa_poets_esotimaote.jl` script in the model repositories.

### Morris sensitivity analysis

Morris sensitivity analysis was used to understand which model parameters were sensitive [42]. The Morris method is a global method that computes an elementary effect value for each parameter by sampling a model performance function, in this case the area under the curve for each model species, over a range of values for each parameter; the mean of elementary effects measures the direct effect of a particular parameter on the performance function, while the variance of each elementary effect indicates whether the effects are non-linear or the result of interactions with other parameters (large variance suggests connectedness or nonlinearity). The Morris sensitivity coefficients were computed using the `DiffEqSensitivity.ji` package [37]. The parameter ranges were established by calculating the minimum and the maximum value for each parameter in the parameter ensemble generated by JuPOETs. Each range was then subdivided into 10,000 samples for the sensitivity calculation. The Morris sensitivity coefficients are calculated using the `compute_sensitivity_coefficients.ji` script in the model repositories.

## Results

### Synthetic circuit architecture

The two genetic circuits (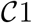 and 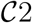) used in this study were based upon the bacterial sigma factor regulatory system (Fig. 1). Sigma factor 70 (*σ*_70_), endogenously present in the extract, was the primary driver of each circuit. In 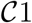, *σ*_70_ induced green fluorescent protein (deGFP) expression was explored in the absence of additional regulators or protein degradation (Fig. 1A). In 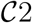, *σ*_70_ induced the expression of sigma factor 28 (*σ*_28_) and deGFP-ssrA (Fig. 1B). Sigma 28 induced the expression of the lambda phage repressor protein cI-ssrA, which was under the *σ*_28_ responsive P28 promoter. The cI-ssrA protein repressed the P70a promoter, thereby down-regulating σ_28_ and deGFP-ssrA transcription [43]. Simultaneously, the C-terminal ssrA degradation tags present on the deGFP and cI proteins were recognized by the endogenous ClpXP protease system in the cell free extract, thereby promoting the degradation of these proteins into peptide fragments [44, 45]. In addition, messenger RNAs (mRNAs) were always subject to degradation due to the presence of degradation enzymes in the extract [45, 46]. Taken together, the interactions of the components manifested in an accumulation of deGFP protein for 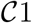, and a pulse signal of deGFP-ssrA in 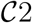. Studying 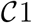 allowed us to estimate parameters governing the interaction of *σ*_70_ with the P70a promoter. Whereas, the 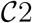 allowed us to characterize the interaction of *σ*_28_ with the P28 promoter, the strength of the transcriptional repression by cI-ssrA, and the kinetics of protein degradation by the endogenous ClpXP protease system. Finally, both circuits tested the effective model formulation for the transcription and translation rates.

**Fig. 1:**
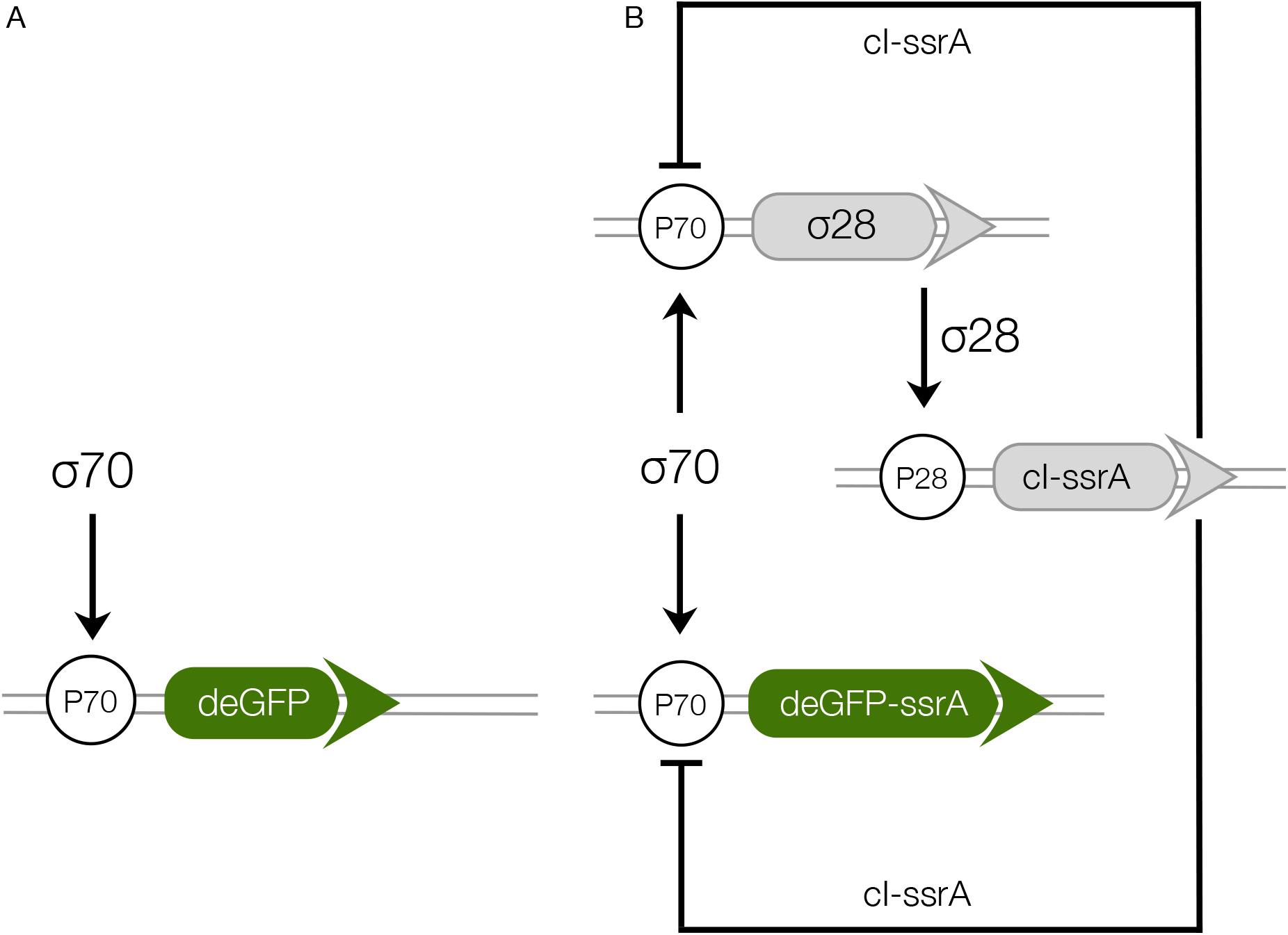
Schematic of the cell free gene expression circuits used in this study. **A**: Sigma factor 70 (*σ*_70_) induced expression of deGFP. **B**: The circuit components encode for a negative feedback loop motif. Sigma factor 28 and deGFP-ssrA genes on the P70a promoters are expressed first because of the endogenous presence of sigma 70 factor in the extract. Sigma factor 28, once expressed, induces the P28a promoter, turning on the expression of the cI-ssrA gene which represses the P70a promoter. The circuit is modified from a previous study [45] by including an ssrA degradation tag on the cI gene.

### Modeling and analysis of the 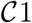 circuit

The effective biophysical transcription and translation model captured σ_70_ induced deGFP expression at the mRNA and protein level within the experimental error for 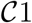 (Fig. 2). JuPOETs produced an ensemble (N = 140) of the 11 unknown model parameters which captured the transcription of mRNA (Fig. 2A) and the translation of deGFP protein (Fig. 2B). The mean and standard deviation of key parameters is given in Table 2. The deGFP mRNA reached its steady state concentration of approximately 580 nM within two hours, and stayed at this level for the remainder of the reaction. Thus, the cell free reaction maintained continuous transcriptional activity with an average mRNA lifetime of 27 minutes; Garamella et al [45] reported a similar lifetime of 17-18 min. On the other hand, deGFP protein concentration increased more slowly, and began to saturate between 8 to 10 hr at approximately 15 *μ*M. Given there was negligible protein degradation (the mean deGFP half-life was estimated to be ~11 days, which was similar to the value of 6 days estimated by Horvath et al, albeit in a different cell free system [33]). The saturating protein concentration suggested that the translational capacity of the cell free system decreased over the course of the reaction. The decrease in translational capacity, which could stem from several sources, was captured in the simulations using a monotonically decreasing translation capacity state variable *ϵ*, and the translational control variable *w* (…). In particular, the mean half-life of translational capacity was estimated to be *τ*_*L*,1/2_ ~ 4 h in the 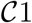 experiments. Taken together, JuPOETs produced an ensemble of model parameters that captured the experimental training data. Next, we considered which 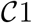 model parameters were important to the model performance using Morris sensitivity analysis, a global sensitivity analysis method.

**Fig. 2:**
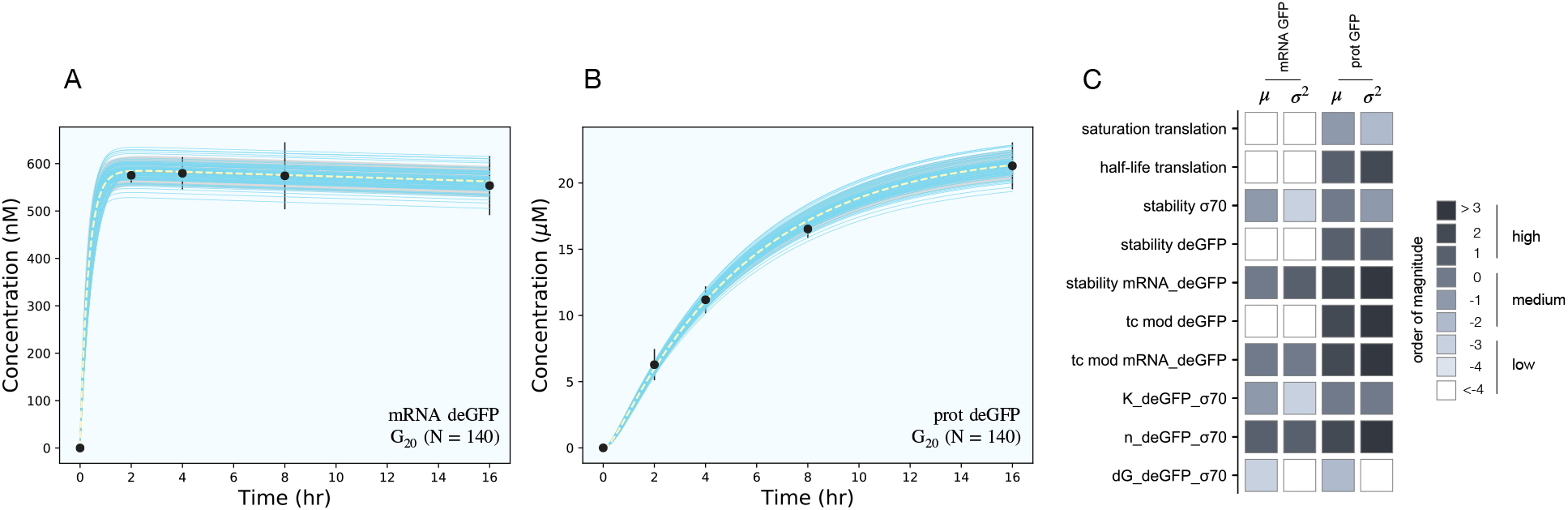
Model simulations versus experimental measurements for *σ*_70_ induced deGFP expression. **A**: Simulated and measured deGFP mRNA concentration versus time using the small circuit G_20_ ensemble (N = 140). **B**: Simulated and measured deGFP protein concentration versus time the small circuit *G*_20_ ensemble (N = 140). **C**: Global sensitivity analysis of the P70-deGFP circuit parameters. Morris sensitivity coefficients were calculated for the unknown model parameters, where the range for each parameter was established from the ensemble. Uncertainty: Simulations and uncertainty quantification are shown for the generation 20 (*G*_20_) ensemble which yielded N = 140 parameter sets that were rank two or below. The parameter ensemble was used to calculate the mean (dashed line) and the 95% confidence estimate of the simulation (gray region). Additionally, the 99% confidence estimate of the mean simulation is shown in orange. Individual parameter set trajectories are shown in blue. Points denote the mean experimental measurement while error bars denote the 95% confidence estimate of the experimental mean computed from at least three replicates.

**Table 2:**
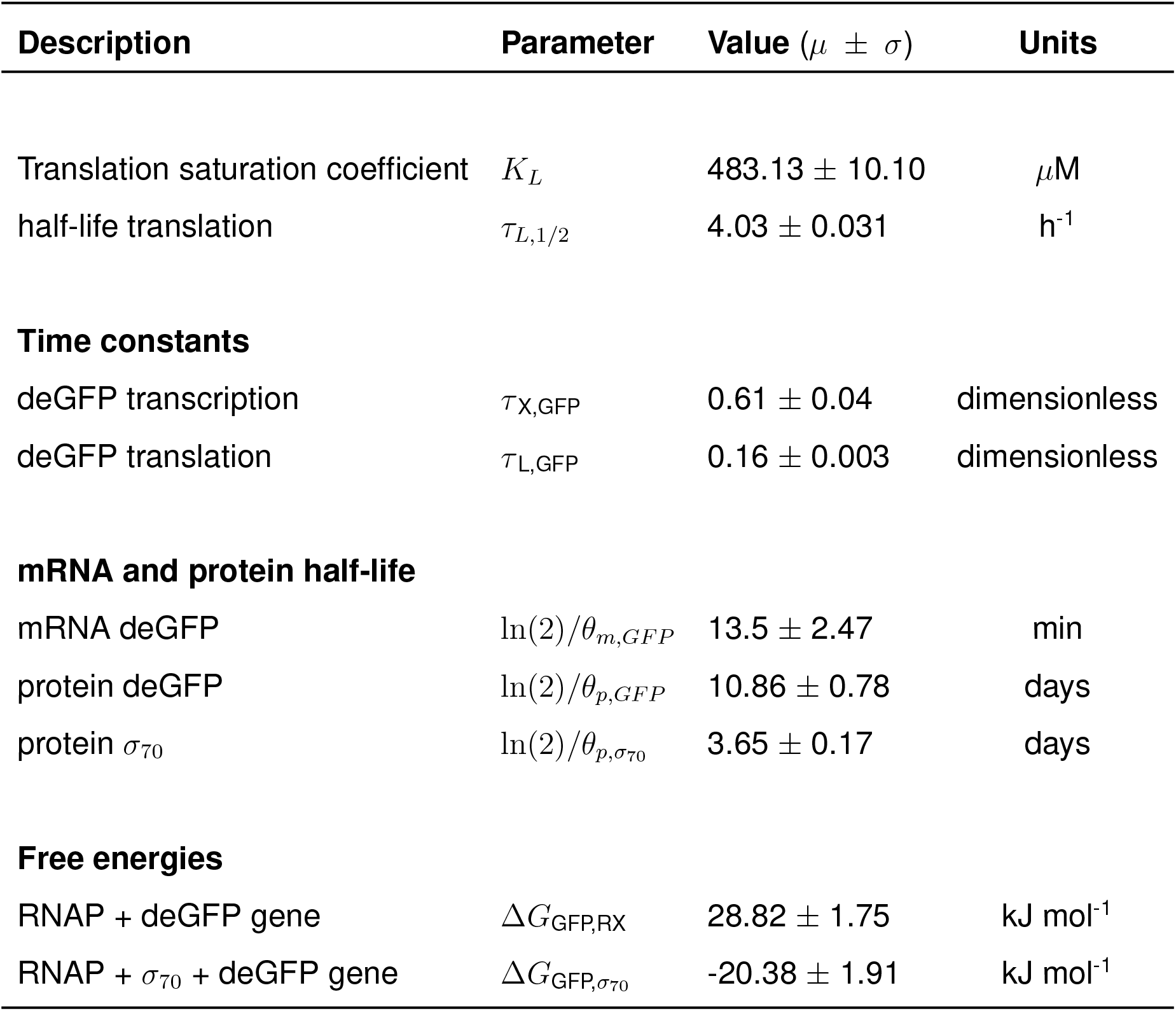
Estimated parameter values for the P70-deGFP model 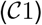. The mean and standard deviation of each parameter value was calculated over the ensemble of parameter sets meeting the rank selection criteria (N = 139).

The importance of 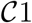 model parameters was quantified using Morris sensitivity analysis (Fig. 2B). The Morris method computes the influence of each parameter, known as the elementary effect, on a model performance function. The mean of elementary effects measures the direct effect of a particular parameter, while the variance indicates whether the effects are non-linear or the result of interactions with other parameters (large variance suggests nonlinearity). The performance function was defined as the area under the curve for each species. The Morris sensitivity measures (mean and variance) were binned into categories based upon their relative magnitudes, from no influence (white) to high influence (black). Only four parameters (translation saturation coefficient *K_L_*, translational capacity half-life *τ*_*L*,1/2_, translation time constant, and protein degradation constant) influenced the protein level. On the other hand, six parameters influenced both mRNA and protein abundance; all six of these parameters were either directly or indirectly associated with transcription. Thus, these parameters influenced the production or stability of mRNA which in turn influenced the protein level. In particular, the mRNA degradation constant and the cooperativity of *σ*_70_ in the P70a promoter function had the largest direct effect and variance. Surprisingly, the ΔG of *σ*_70_/RNAP/promoter configuration was the least influential of the six parameters and had a small elementary effect variance. Taken together, Morris sensitivity analysis of the 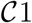 model parameters highlighted the hierarchical structure of the transcriptional and translation model, suggesting experimentally tunable parameters such as mRNA stability were globally important. Next, we used the ensemble of P70a, time constant and degradation parameters estimated for 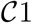 to constrain the analysis of 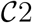.

### Modeling and analysis of the 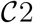 circuit

The effective biophysical transcription and translation model captured the deGFP-ssrA expression dynamics in the negative feedback circuit 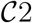 (Fig. 3A). JuPOETs produced an ensemble (N = 498) of the 33 unknown model parameters which captured transcription and translation dynamics for *σ*_28_, cI-ssrA and deGFP-ssrA. The mean and standard deviation of key parameters is given in Table 3. Compared with the estimated parameters for 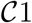, the 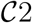 model had almost a two fold change in the half life of translation and the translation saturation coefficient. Similarly, there were variations in the values of the transcription and translation time constants for the two systems. However, for both circuits, the small values of the transcription and translation time constants qualitatively suggested elongation limited reactions; the exception was *σ*_28_ translation which was closer to initiation limited. Unlike 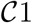, the mRNA expression pattern for *σ*_28_ and deGFP-ssrA both showed an initial spike, to a concentration similar with the previous pseudo steady state, before the cI-ssrA regulator protein could be expressed. However, once cI-ssrA began to accumulate, the concentrations of the regulated mRNAs dropped by approximately an order of magnitude compared with the unregulated case. Again, as shown with 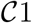, the regulated mRNA concentrations reached an approximate steady-state. This further confirmed continuous transcription and mRNA degradation throughout the cell free reaction. The mean estimated mRNA lifetime for cI-ssrA and deGFP were similar (approximately 16 min), while the degradation of *σ*_28_ mRNA was predicted to be slower (mean mRNA lifetime was estimated to be approximately 30 min). Lastly, the mean peak degradation rate for GFP was approximately 47 nM/min, while the mean peak cI-ssrA degradation rate was predicted to be approximately 63 nM/min; both of these degradation rate estimates were consistent with the previously reported range of 15 nM/min - 150 nM/min measured by Garamella et al [45].

**Fig. 3:**
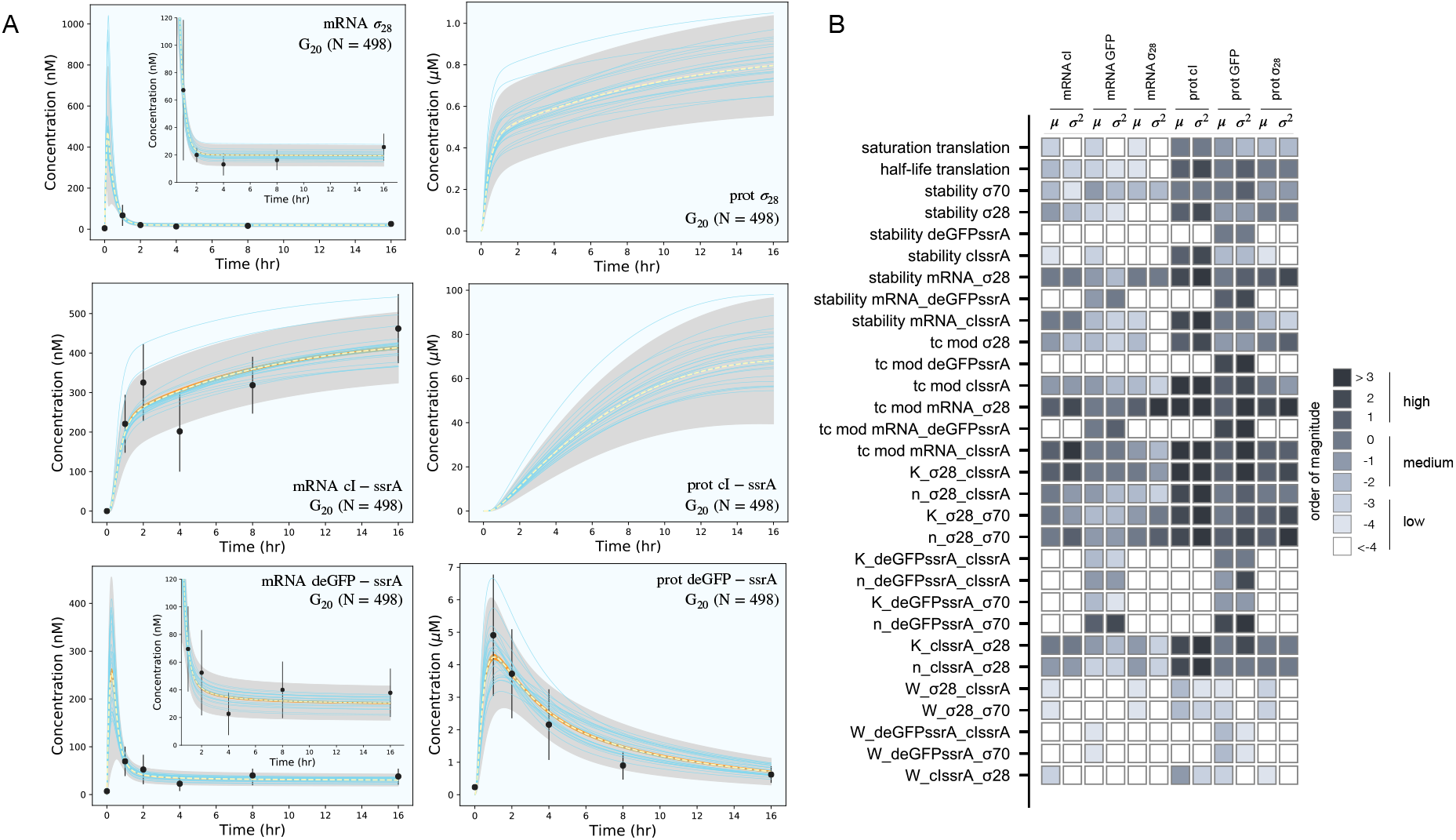
Model simulations versus experimental measurements for the negative feedback deGFP-ssrA circuit. **A**: Model simulations of the negative feedback deGFP-ssrA circuit using the *G*_20_ ensemble (N = 498). Uncertainty: Simulations and uncertainty quantification are shown for the generation 20 (*G*_20_) ensemble which yielded N = 489 parameter sets (rank two or below). The parameter ensemble was used to calculate the mean (dashed line) and the 99% confidence estimate of the simulation (gray region). Additionally, the 99% confidence estimate of the mean simulation is shown in orange. Individual parameter set trajectories are also shown in blue. Points denote the mean experimental measurement while error bars denote the 95% confidence estimate of the experimental mean computed from at least three replicates. **B**: Global sensitivity analysis of the negative feedback deGFP-ssrA circuit parameters. Morris sensitivity coefficients were calculated for the unknown model parameters, where the range for each parameter was established from the ensemble.

**Table 3:**
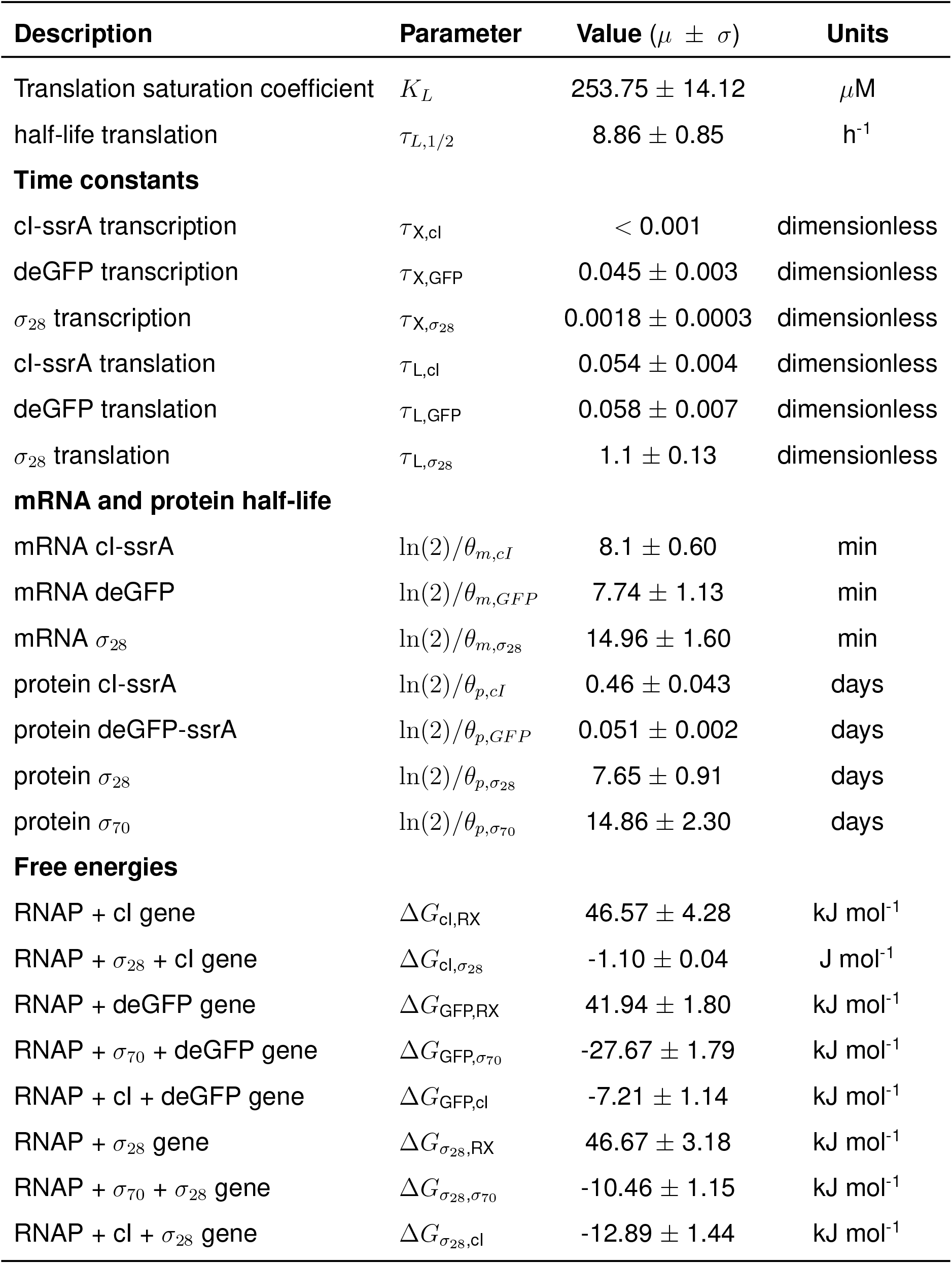
Estimated parameter values for the negative feedback circuit 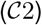. The mean and standard deviation for each parameter was calculated over the ensemble of parameter sets (N = 498).

| Description | Parameter | Value ($\mu \pm \sigma$) | Units |
| --- | --- | --- | --- |
| Translation saturation coefficient | $K_L$ | $253.75 \pm 14.12$ | $\mu\text{M}$ |
| half-life translation | $\tau_{L,1/2}$ | $8.86 \pm 0.85$ | $\text{h}^{-1}$ |
| **Time constants** |  |  |  |
| cl-ssrA transcription | $\tau_{X,\text{cl}}$ | $< 0.001$ | dimensionless |
| deGFP transcription | $\tau_{X,\text{GFP}}$ | $0.045 \pm 0.003$ | dimensionless |
| $\sigma_{28}$ transcription | $\tau_{X,\sigma_{28}}$ | $0.0018 \pm 0.0003$ | dimensionless |
| cl-ssrA translation | $\tau_{L,\text{cl}}$ | $0.054 \pm 0.004$ | dimensionless |
| deGFP translation | $\tau_{L,\text{GFP}}$ | $0.058 \pm 0.007$ | dimensionless |
| $\sigma_{28}$ translation | $\tau_{L,\sigma_{28}}$ | $1.1 \pm 0.13$ | dimensionless |
| **mRNA and protein half-life** |  |  |  |
| mRNA cl-ssrA | $\ln(2)/\theta_{m,\text{cl}}$ | $8.1 \pm 0.60$ | min |
| mRNA deGFP | $\ln(2)/\theta_{m,\text{GFP}}$ | $7.74 \pm 1.13$ | min |
| mRNA $\sigma_{28}$ | $\ln(2)/\theta_{m,\sigma_{28}}$ | $14.96 \pm 1.60$ | min |
| protein cl-ssrA | $\ln(2)/\theta_{p,\text{cl}}$ | $0.46 \pm 0.043$ | days |
| protein deGFP-ssrA | $\ln(2)/\theta_{p,\text{GFP}}$ | $0.051 \pm 0.002$ | days |
| protein $\sigma_{28}$ | $\ln(2)/\theta_{p,\sigma_{28}}$ | $7.65 \pm 0.91$ | days |
| protein $\sigma_{70}$ | $\ln(2)/\theta_{p,\sigma_{70}}$ | $14.86 \pm 2.30$ | days |
| **Free energies** |  |  |  |
| RNAP + cl gene | $\Delta G_{\text{cl,RX}}$ | $46.57 \pm 4.28$ | $\text{kJ mol}^{-1}$ |
| RNAP + $\sigma_{28}$ + cl gene | $\Delta G_{\text{cl},\sigma_{28}}$ | $-1.10 \pm 0.04$ | $\text{J mol}^{-1}$ |
| RNAP + deGFP gene | $\Delta G_{\text{GFP,RX}}$ | $41.94 \pm 1.80$ | $\text{kJ mol}^{-1}$ |
| RNAP + $\sigma_{70}$ + deGFP gene | $\Delta G_{\text{GFP},\sigma_{70}}$ | $-27.67 \pm 1.79$ | $\text{kJ mol}^{-1}$ |
| RNAP + cl + deGFP gene | $\Delta G_{\text{GFP,cl}}$ | $-7.21 \pm 1.14$ | $\text{kJ mol}^{-1}$ |
| RNAP + $\sigma_{28}$ gene | $\Delta G_{\sigma_{28},\text{RX}}$ | $46.67 \pm 3.18$ | $\text{kJ mol}^{-1}$ |
| RNAP + $\sigma_{70}$ + $\sigma_{28}$ gene | $\Delta G_{\sigma_{28},\sigma_{70}}$ | $-10.46 \pm 1.15$ | $\text{kJ mol}^{-1}$ |
| RNAP + cl + $\sigma_{28}$ gene | $\Delta G_{\sigma_{28},\text{cl}}$ | $-12.89 \pm 1.44$ | $\text{kJ mol}^{-1}$ |

The secondary effect of cI-ssrA repression was visible in the cI-ssrA mRNA expression pattern. The expression of cI-ssrA was induced by *σ*_28_, however, *σ*_28_ expression was repressed by cI-ssrA, thereby completing a negative feedback loop. Initially, before appreciable levels of cI-ssrA had been translated, the cI-ssrA transcription rate was maximum (approximately 200 nM/h). However, the transcription rate decreased to approximately 12 nM/h after two hours and remained constant for the remainder of the cell free reaction. Similarly, transcription rates for *σ*_28_ (approximately 1200 nM/h) and deGFP-ssrA (approximately 750 nM/h) were initially at a maximum due to the presence of endogenous *σ*_70_, but then quickly dropped as cI protein accumulated. Protein synthesis followed a similar trend, with the translation rates for *σ*_28_ and deGFP-ssrA initially present at their maximum values before quickly dropping. After one hour, deGFP levels reached a peak and decayed due to the ClpXP-mediated degradation, whereas *σ*_28_ protein levels continued to slowly rise at a steady rate (approximately 15 nM/h). The 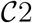 model also predicted the expected lag present during the initial phase of cI-ssrA protein synthesis due to the need for *σ*_28_ protein to reach appreciable levels. Moreover, the combination of high cI-ssrA mRNA abundance (expressed because *σ*_28_ does not have a degradation tag) and ClpXP-mediated degradation led to the saturation of the cI-ssrA protein concentration. However, the cI-ssrA protein concentration could not be verified because we did not have an experimental measurement for this species. Taken together, the effective model simulated cell free expression dynamics for 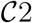. Next, we considered which 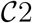 model parameters were important using Morris sensitivity analysis.

Morris sensitivity analysis of the negative feedback circuit 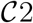 stratified the parameters into locally and globally important groups (Fig. 3B). The influence of 33 parameters was computed using the area under the curve of each mRNA and protein species as the performance function. The Morris sensitivity metrics (mean and variance) were binned into categories based upon their relative magnitudes, from no influence (white) to high influence (black). Some parameters affected only their respective mRNA or protein target, whereas others had widespread effects. For example, the time constant (tc) modifiers, stability of deGFP-ssrA protein and mRNA, and the binding dissociation constant (K) and cooperativity parameter (n) of cI-ssrA and *σ*_70_ for the deGFP-ssrA promoter affected only the values of deGFP-ssrA protein and mRNA. On the other hand, the tc, stability, K and n parameters for *σ*_70_, *σ*_28_, or cI-ssrA influenced mRNA and protein expression globally. The *σ*_70_ and *σ*_28_ proteins acted as inducers or repressors for more than one gene product: *σ*_70_ induced both deGFP-ssrA and *σ*_28_, and cI-ssrA protein repressed both of these genes. Degradation constants (denoted as stability) affected the half-lives of the transcribed messages or the translated proteins in the mixture, while the time constant modifiers changed the time required to form the open gene complex (or translationally active complex). Dissociation and cooperativity constants affected the binding interactions of the inducer (or repressor in the case of cI-ssrA) in the promoter control function. Varying these parameters, therefore, had a strong effect on their respective targets. Similarly, the translation saturation and its half-life, which captured the depletion in the translation activity over the course of the reaction, not only affected protein levels but also mRNA levels. This is because these parameters tuned the rate of formation of cI-ssrA, which in turn affected the mRNA levels of its gene targets. Given that cI-ssrA was the main regulator (repressor) of the circuit, the parameters that dictated the levels of cI-ssrA mRNA and protein had a global effect. We also observed high sensitivity variance for several parameters, in particular involving cI-ssrA. For example, the time constant modifiers for cI-ssrA mRNA and protein had a two-pronged effect. On the one hand, they positively influenced the transcription/translation rates of the gene and mRNA products, directly increasing the cI-ssrA protein. On the other hand, increased cI-ssrA expression reduced the level of *σ*_28_, in turn reducing the cI-ssrA levels over time. Taken together, Morris sensitivity analysis of the 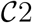 model stratified that parameters into local and globally important groups, with the parameters governing the synthesis rates of the cI-ssrA mRNA and protein being globally important. The sensitivity analysis also gave insight into the organization of the circuit, suggesting cI to be highly connected within the circuit.

## Discussion

In this study, we developed an effective biophysical modeling approach to simulate transcription (TX) and translation (TL) processes occurring in a cell free system. We tested this approach by simulating the dynamics of two cell free synthetic circuits (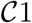 and 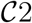). The model formulation, and parameter values were mechanistic and largely derived from literature. For example, characteristic values for *τ_X_* and *K_X_*, the time and saturation constants for transcription, were approximated from *in vitro* experiments using an abortive initiation assay [47]. The RNAP and ribosome elongation rates were taken from Garamella et al. [45], while other parameter were estimated from BioNumbers [48]. Similarly, the weights appearing in the transcription control function *u*(…) were based upon the Gibbs energies of the respective promoter configurations, while the form of the transcriptional control functions was derived from a statistical mechanical treatment of promoter activity [38–40]. However, there were parameters that were not available from literature; in these cases multiobjective optimization was used to estimate these parameters from mRNA and protein measurements. For 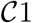, sigma factor 70 (*σ*_70_) induced expression of green fluorescent protein (deGFP), the time constants, degradation rates, and other parameters governing deGFP expression were estimated from measurements of deGFP mRNA and protein. These estimates were then used to constrain the parameter search for 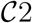, which involved deGFP expression subject to negative feedback and programmed protein degradation. We estimated which model parameters were important to the performance of 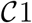 and 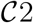 using Morris sensitivity analysis. Sensitivity analysis results for 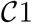 were expected; the time constant for transcription, the stability of the deGFP message and the cooperativity of *σ*_70_ were all important parameters. On the other hand, the sensitivity analysis results for 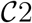 were more nuanced, with parameters (and associated species) being stratified into locally and globally important groups; the performance of 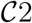 was most sensitive to the parameters controlling the cI-ssrA mRNA and protein abundance.

The effective TX/TL modeling approach described here has several potential applications. For example, a challenge of *in vivo* constraint based modeling is the description of gene expression [49]. Boolean and probabilisitic approaches [50–52] have been developed to address this challenge. However, the transcriptional state of a boolean gene is either on or off based on the state of its regulators, thus, gene expression is course grained. The current modeling approach could be an interesting mechanistic alternative to the boolean framework that utilizes a continuous description of gene expression dynamics and transcriptional regulation. In particular, the rules encoding typical boolean gene expression networks are easily translatable into the rational promoter functions described here, however, the estimation of the parameters appearing in these promoter functions, especially in an *in vivo* context, remains an open question. Another interesting application could be a description of the integration of gene expression with metabolism in cell free synthetic biology applications. For example, synthetic biology studies often neglect the role that metabolism plays in the expression of synthetic circuits. Ultimately, metabolism is centrally important to the operation of any synthetic circuit; gene expression is strongly dependent upon the metabolic resources provided by catabolic processes. We have recently started to explore this question by integrating effective transcription and translation models with metabolic networks in cell free reactions e.g., [33, 53], and also developing experimental tools to measure metabolite concentrations in cell free systems [54]. However, these previous transcriptional and translations models (and similar precursor models simulating eukaryotic processes [55, 56]) were less developed than those presented here. Taken together, the effective modeling approach described here can potentially be used to simulate transcription and translation processes in a variety of applications.

The effective TX/TL model described the experimental mRNA and protein training data. However, there were several important questions to be addressed by future studies. First, the model formulation described the data, but did not predict dynamics outside of the training set. If this approach is to be useful to the synthetic biology community, or more broadly as an effective biophysical technique to model *in vivo* gene expression dynamics for applications such as regulatory flux balance analysis, we need to have confidence that the modeling approach is predictive. Thus, while we have established a descriptive model, we have to yet to establish a predictive model. Next, there were several technical or mechanistic questions that should be explored further. For example, cI-ssrA represses the activity of the P70a promoter via interaction with its OR2 and OR1 operator sites; in this study we considered only a single operator site suggesting that we potentially underestimated the potency of cI repression in the deGFP and *σ*_28_ promoter functions, see the multiplication rule [57]. Further, we used a first order approximation of ClpXP mediated protein degradation, while Garamella et al. [45] described this degradation as zero order. Similarly, we did not establish the concentration of ClpXP in the commercially available cell free reaction mixture. The levels of this protein complex could be an important factor controlling protein degradation. Next, we should compare the current modeling approach, and the values estimated for the model parameters, with the study of Marshall and Noireaux [32]. For example, one of the potential limitations of the current study (that was addressed by Marshall and Noireaux [32]) is that we did not consider a separate species for dark GFP. In our previous RNA circuit modeling [58], we did include this term, but failed to do so here. We expect inclusion of a dark versus light GFP species could influence the values for the estimated parameters, particularly the translation time constants. However, previous reports suggested the *in vitro* maturation time of deGFP was approximately 8 min [59], much faster than the typical maturation times for GFP of one hour *in vivo* [60, 61]. Thus, the impact of including a dark versus light GFP species may not be worth the increased model complexity. Lastly, we should validate the values estimated for the binding function parameters and the promoter configuration free energies. Maeda et al measured the binding affinities of the seven *E. coli σ* factors with RNAP [62]; while not directly comparable, these measurements give an order of magnitude characteristic value for the dissociation constants appearing in the promoter binding functions. Further, there have been several studies that have quantified the binding energies of promoter configurations e.g., [38, 63–65]. A perfunctory inspection of the values estimated in this study suggested our estimated free energy values, while the same order of magnitude as previous studies in many cases, did have values that were off by a factor of up to an order of magnitude compared with literature (albeit for different promoters). In particular, the positive Gibbs energy estimated for free RNAP binding leading to transcription was likely too large, while the magnitude of other values such as the energy of cI repression of *σ*_28_ expression was likely too small. Thus, these other studies could serve as a basis to validate our estimates, and perhaps more importantly constrain the parameter search space for future studies.

## Author Contributions

J.V directed the study. M.V, S.V, H.L and A.A conducted the cell free experimental measurements. J.V, M.V and A.A developed the reduced order models and the parameter ensemble. M.V, A.A and J.V analyzed the model ensemble, and generated figures for the manuscript. The manuscript was prepared and edited for publication by A.A, M.V, S.V and J.V. All authors reviewed this manuscript.

## Acknowledgments

We gratefully acknowledge the suggestions from the anonymous reviewers to improve this manuscript.

## Data Availability Statement

Model code is available under an MIT software license from the Varnerlab GitHub repository [34]. The mRNA and protein measurements presented in this study are available in the data directory of the model repositories in comma separated value (CSV) and Microsoft Excel format.

